# Tau assemblies do not behave like independently acting prion-like particles in mouse neural tissue

**DOI:** 10.1101/2021.01.13.426340

**Authors:** Aamir S. Mukadam, Lauren V. C. Miller, Claire S. Durrant, Marina J. Vaysburd, Taxiarchis Katsinelos, Benjamin J. Tuck, Sophie Sanford, Olivia Sheppard, Claire Knox, Shi Cheng, Leo C. James, Michael P. Coleman, William A. McEwan

## Abstract

A fundamental property of infectious agents is their particulate nature: infectivity arises from independently-acting particles rather than as a result of collective action. Assemblies of the protein tau can exhibit seeding behaviour, potentially underlying the apparent spread of tau aggregation in many neurodegenerative diseases. Here we ask whether tau assemblies share with classical pathogens the characteristic of particulate behaviour. We used organotypic hippocampal slice cultures from P301S tau transgenic mice in order to precisely control the concentration of extracellular tau assemblies. Whilst untreated slices displayed no overt signs of pathology, exposure to tau assemblies could result in the formation of intraneuronal, hyperphosphorylated tau structures. However, seeding ability of tau assemblies did not titrate in a one-hit manner in neural tissue. The results suggest that seeding behaviour of tau only arises at supra-physiological concentrations, with implications for the interpretation of high-dose intracranial challenge experiments and the possible contribution of seeded aggregation to human disease.

## Introduction

Neurodegenerative diseases are typified by the accumulation of specific proteins into fibrillar assemblies. In around twenty distinct neurodegenerative diseases, including the most common, Alzheimer’s disease, the protein tau forms hyperphosphorylated, filamentous inclusions within the cytoplasm of neurons. Evidence from human genetics suggests that tau accumulation can be a direct cause of neurodegeneration since around 50 distinct mutations in *MAPT*, the gene that encodes tau, cause inherited forms of dementia with evidence of tau filaments [1]. The origin of tau assemblies in the human brain remains uncertain. Cell-autonomous processes may lead to the spontaneous nucleation of oligomeric forms of tau within the cytoplasm of neurons. Some of these assemblies adopt filamentous conformations that are able to undergo extension by the addition of tau monomers to the filament ends. Over the past decade it has been postulated that, in addition to these cell-autonomous mechanisms, tau pathology may occur through a spreading or prion-like mechanism [2]. Several lines of evidence demonstrate that assemblies of tau can be taken up into cells, whereupon they seed the conversion of native tau to the assembled state. Addition of tau assemblies to the exterior of cells, or the injection of tau assemblies to the brains of tau-transgenic mice, can induce intracellular tau assembly in the recipient [3–5].

Population cross-sectional studies demonstrate that tau pathology follows a predictable pattern over time and space in the human brain consistent with spreading, potentially via a prion-like mechanism. Immunoreactivity to antibodies such as AT8, which detects tau that is abnormally phosphorylated at positions S202 and T205 [6], progresses in a manner that can be systematically categorised into stages according to anatomical distribution (Braak stages 0 - VI) [7,8]. In young adults, some AT8 immunoreactivity is observed in the vast majority of brains by the third decade of life. However, it is generally confined to neurons within the locus coeruleus (LC) in the brainstem (Braak pretangle stages 0 a-c and 1a,b). Subsequently, AT8 staining is observed in the entorhinal cortex (EC) and hippocampus (HC) (Braak stages I-II). Later stages are characterised by progressive dissemination and increasing density of staining in neocortical regions (Braak stages III-VI). These late stages are associated with severe disease and the overall burden of tau pathology negatively corelates with cognitive function [9].

Though intracranial challenge experiments demonstrate that seeded aggregation can in principle occur, they provide little insight as to whether physiological concentrations of extracellular tau species might support prion-like activity. The concentration of tau in wildtype mouse interstitial fluid (ISF) is around 50 ng/ml total tau (equivalent to ∼1 nM tau monomer). Mouse ISF levels typically exceed cerebrospinal fluid (CSF) tau levels by around 10-fold [10]. In humans between ages 21 to 50 years, CSF total tau is below 300 pg/mL increasing to 500 pg/mL over age 70 [11] – approximately 7 to 12 pM if considering the average mass of full length tau isoforms. Levels are increased 2-3 fold in Alzheimer’s disease [12]. If a similar relationship between ISF and CSF tau concentration exists in humans as in mice, ISF tau levels are likely in the order of 100 pM, rising to 300 pM in Alzheimer’s disease. Intracranial injection experiments typically supply tau in the high micromolar range. Even if this were distributed broadly across the brain, micromolar concentrations would be exceeded and local concentration at the injection site may plausibly be 100-fold greater. Thus, intracranial injection experiments likely exceed physiological concentrations of extracellular tau by two to seven orders of magnitude.

For classical infectious agents, infectivity is related to dose by a “one-hit” relationship wherein the amount of infectivity decreases linearly upon dilution until end-point [13]. This property is also evident in PrP^Sc^ prions, though it is complicated by the presence of multiple aggregation states and the size distribution of particles [14]. The relationship between dose and prion-like activity for tau has not been established. It is therefore currently not possible to reconcile high-dose challenge experiments with the low concentrations of tau observed in the extracellular spaces of the brain. To address this, we developed a model of seeded tau aggregation in mouse organotypic hippocampal slice cultures, allowing direct control of the concentration of tau neurons were exposed to. Brain slice cultures have been used for 40 years [15], though developments in recent years have rendered them increasingly relevant for the study of neurodegenerative diseases [16–20]. We prepared slices from transgenic mice with the *MAPT* P301S mutation [21], which is causative of fronto-temporal dementia and displays accelerated fibrilisation compared to wildtype tau [22]. ISF concentrations of tau in P301S transgenic mice have previously been measured at about 5 times that of wildtype animals at around 5 nM monomer equivalent, versus 1 nM in wildtype, consistent with the reported 5-fold over-expression of tau [10].

Using our system, which relies on physiological neuronal uptake of tau aggregates supplied to the media, we show that neurons within CA1 are preferentially susceptible to seeded aggregation, displaying intracellular hyperphosphorylated tau tangles. We find that seeding activity cannot be titrated down and only occurs at high concentrations of tau assemblies. Crucially, at between 30 and 100 nM, the concentrations of tau assemblies required to initiate seeding exceed reported measures of physiological ISF and CSF tau. Our results imply that a model of tau spread via seeded aggregation requires these concentrations to be locally exceeded or requires other mechanisms not captured here to facilitate seeded aggregation.

## Results

Sagittal hippocampal slices of ∼300 μm thickness were prepared from homozygous P301S tau-transgenic mice at age 7 d. Hippocampal structures are well developed at this age, yet the tissue exhibits plasticity that aids recovery from the slicing procedure [23]. We stained OHSCs with antibodies against markers of the major cell types of the brain: neurons (Map2), microglia (Iba1) and astrocytes (Gfap) [Figure 1a]. Similar to previous studies [19], neurons were found to maintain extensive arborisation with evidence of intact neuronal tracts. Microglia were observed with normal morphology with extensive processes, similar to quiescent cells in whole brains [24]. Astrocytes were also well represented throughout the cultures. Immunostaining for tau revealed widespread expression in all regions of the hippocampus [Figure 1b]. After 5 weeks in culture no overt signs of tau pathology were apparent, as visualised by staining with AT8 [Figure 1c]. These results demonstrate that hippocampal architecture and cell types from P301S tau transgenic mice are maintained through the slicing and culture process. Importantly, they demonstrate that OHSCs from P301S tau transgenic mice do not undergo detectable spontaneous aggregation over this time period.

**Figure 1:**
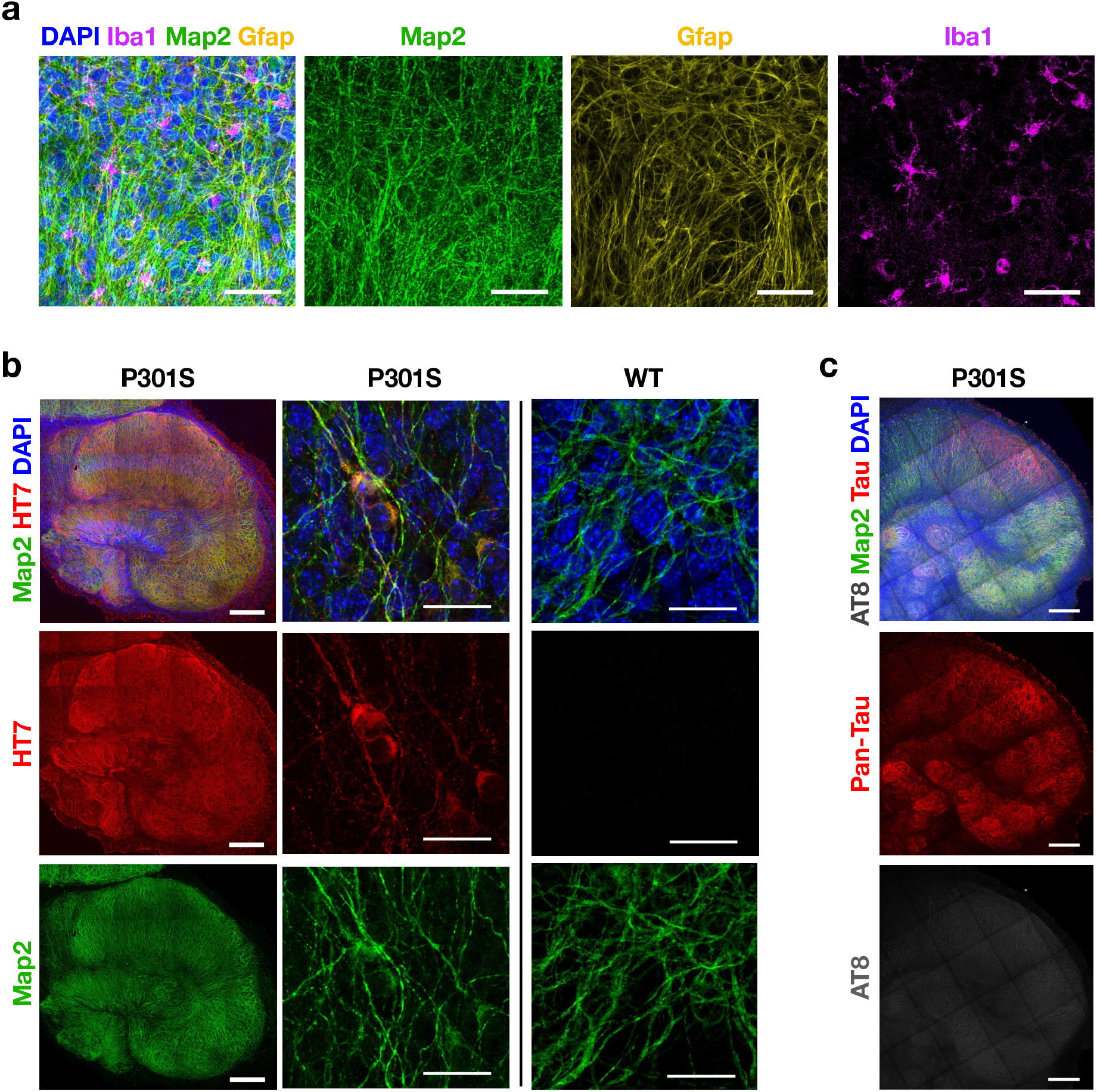
OHSCs maintain cellular diversity and display no spontaneous tau pathology. **a**. OHSCs from mice transgenic for P301S tau were fixed after 2 weeks in culture and stained for nuclei (DAPI), the neuronal marker Map2, the astrocyte marker Gfap, and the microglial marker Iba1. Scale bars are 50 µm. **b**. OHSCs from P301S tau transgenic mice are positive for human tau-specific antibody HT7 whereas OHSCs from WT mice are not. Scale bars 250 µm and 25 µm. **c**. OHSCs from P301S tau transgenic mice after 5 weeks in culture display only background levels of staining with the phospho-tau specific antibody AT8. Slices were stained with DAPI and Map2 as above and with pan-tau. Scale bars are 250 µm.

To investigate the response of slice cultures to challenge with tau assemblies, we prepared tau from two independent sources. First, we expressed the 0N4R isoform of tau bearing the P301S mutation in *E. coli*. Recombinant protein was incubated with heparin and, following a lag period, was found to give a fluorescence signal in the presence of thioflavin T, a dye whose fluorescence increases upon binding to β-sheet rich amyloid structures [Figure 2a]. Negative stain transmission electron microscopy revealed the presence of abundant filamentous structures [Figure 2b]. Second, we prepared the sarkosyl-insoluble (SI) fraction from aged P301S tau transgenic mice, a procedure that enriches insoluble tau species. Brain-derived assemblies were subjected to western blot, confirming the presence of hyperphosphorylated, insoluble tau [Figure 2c]. The samples were quantified using a dot-blot method using recombinant fibrillar tau as a standard [Supplementary Figure 1]. Tau assemblies were added to HEK293 cells stably expressing 0N4R P301S tau-venus, a reporter cell line for seeded aggregation [25]. In this assay, transfection reagents are used to deliver tau assemblies into cells, whereupon tau-venus is observed to form puncta over 1-2 d. This aggregation was previously found to result in the accumulation of tau-venus in the sarkosyl-insoluble pellet [25]. In the present study, abundant venus-positive puncta were detected following challenge with recombinant fibrils or mouse brain derived tau [Figure 2d]. To investigate whether these seeded assemblies bore markers of tau hyperphosphorylation, we stained with the monoclonal antibody AT8 and AT100, which recognises tau phosphorylated at pT212 and pS214. These epitopes occur on tau filaments extracted from post-mortem tauopathy brains including those from Alzheimer’s disease patients. We observed that challenge with recombinant tau assemblies resulted in colocalization between tau-venus puncta and AT8 or AT100 [Figure 2e,f]. We therefore concluded that our tau preparations contained species able to induce *bona fide* seeded aggregation in recipient cells.

**Figure 2:**
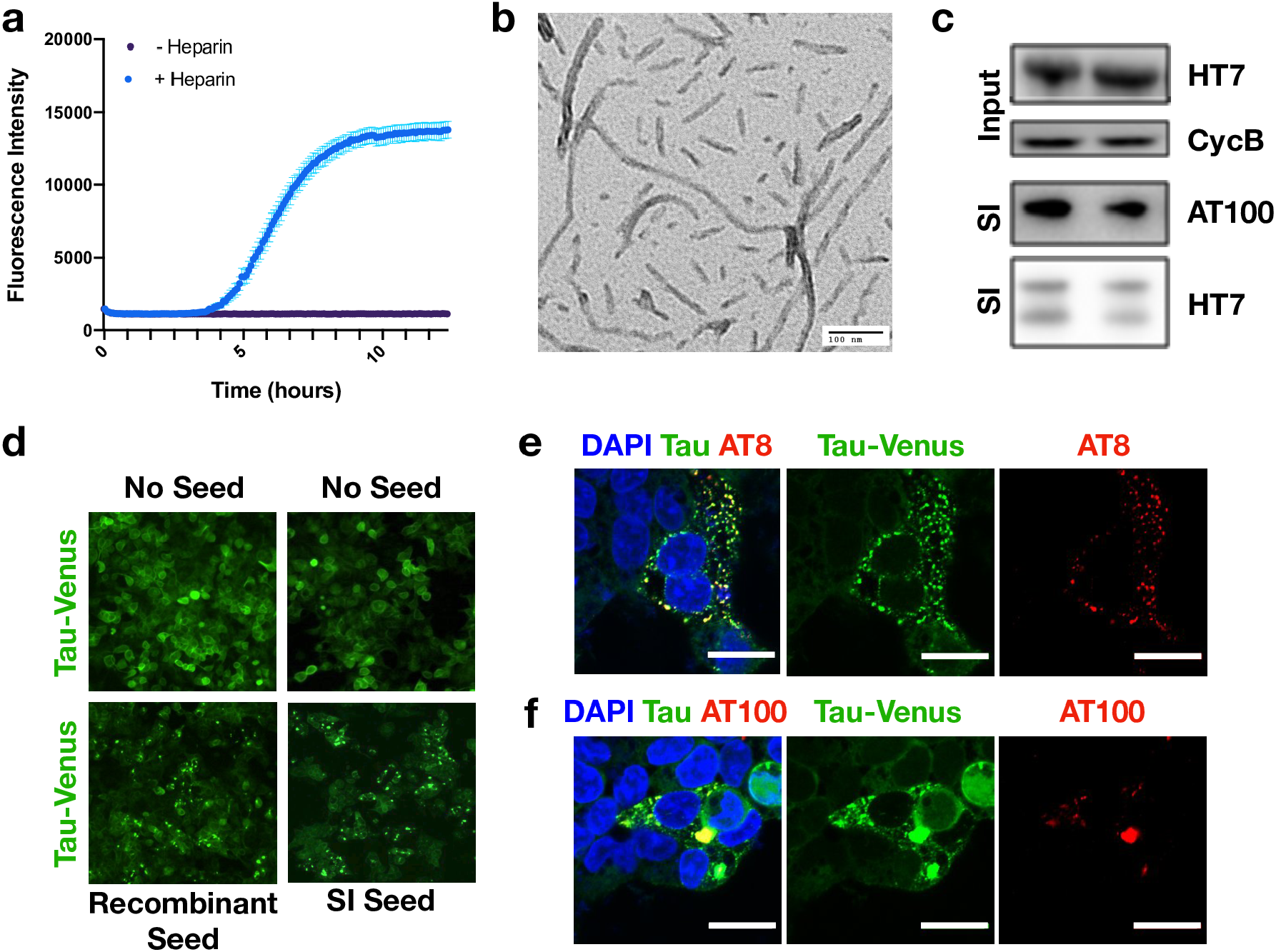
Characterisation of tau assemblies. **a**. Aggregation kinetics for recombinantly produced P301S tau, monitored by ThT fluorescence. **b**. Representative TEM image of recombinantly produced P301S tau assemblies, aggregated with heparin. **c**. Aged P301S tau transgenic mouse brain homogenate was immunoblotted for human tau (HT7 antibody) to detect P301S tau and with Cyclophilin B which served as a loading control. Presence of SI tau was confirmed with HT7 (total tau) and AT100 (tau phosphorylated at pT212, pT214). Lanes represent homogenate and SI fractions from different mice which were subsequently pooled. **d**. Representative images from the tau-venus seeding assay 48 h after challenge with either recombinant P301S tau assemblies or SI tau in the presence of LF2000. **e**,**f**. Tau-venus aggregates observed following challenge with tau assemblies stain with AT8 and AT100 demonstrating that the induced tau aggregates are phosphorylated. Scale bars are 20 µm.

Next, we challenged OHSCs with tau assemblies. Recombinant or mouse brain-derived tau assemblies were supplied to the culture media for a period of three days followed by twice-weekly media changes [Figure 3a]. Three weeks after challenge with 100 nM recombinant tau assemblies or 5 μl SI tau we observed pronounced AT8 staining [Figure 3b][Supplementary Figure 1], suggesting the presence of mature hyperphosphorylated tau assemblies. The addition of monomeric tau did not induce these same structures, indicating that the misfolded state of tau was responsible for seeded aggregation [Figure 3b]. Furthermore, addition of tau assemblies to OHSCs prepared from wildtype mice did not induce seeded aggregation suggesting that the transgenic P301S tau construct is responsible for the phenotype [Figure 3b]. The levels of AT8 between unseeded WT and P301S OHSCs were non-significant [Figure 3c]. We also observed the accumulation of sarkosyl-insoluble species following seeding in P301S transgenic OHSC but not wildtype OHSCs [Figure 3d,e][Supplementary Figure 2]. Taken together, these data demonstrate that insoluble, hyperphosphorylated tau assemblies can be induced in transgenic OHSCs by the addition of exogenous tau assemblies.

**Figure 3:**
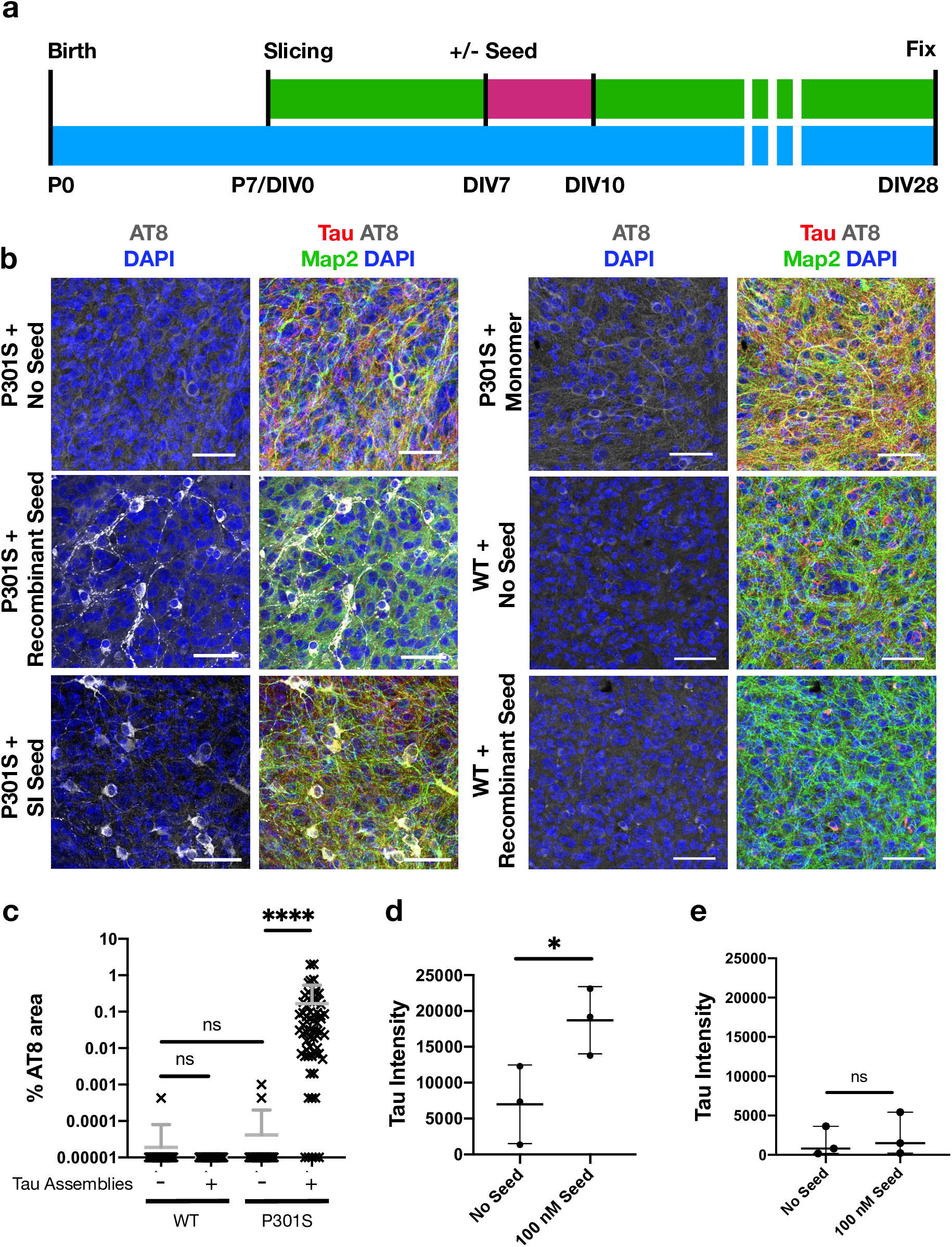
Challenging OHSCs with exogenous tau assemblies induces seeded tau aggregation. **a**. Schematic of OHSC preparation and treatment. Slices were prepared from P7 pups. Tau assemblies were added to the media and incubated for 72 h. A complete media change was carried out at the end of the seeding period (pink). At other times (green) 50% media changes were performed twice weekly until fixation at 28 days *in vitro* (DIV). **b**. P301S OHSCs were challenged with either 100 Nm recombinant tau assemblies, 100 nM monomeric tau, 5 µL of SI tau or buffer only. WT OHSCs were challenged with 100 nM recombinant tau assemblies or buffer only. Scale bars are 50 µm. **c**. Quantification of seeding levels in WT and P301S OHSCs, upon the addition of 100 nM recombinant tau assemblies or buffer only. Statistical significance determined by Kruskal-Wallis Test by ranks and Dunn’s multiple comparisons test (Slices from 3 different mice, per condition. **** P<0.0001). **d**. Analysis of the SI fraction of P301S OHSCs with and without the addition of 100 nM recombinant tau assemblies. **e**. Analysis of the SI fraction of WT OHSCs with and without the addition of 100 nM recombinant tau assemblies. Data normalised to overall levels of tau and loading control. Statistical significance determined by unpaired t-test with Welch’s correction (Slices from 3 different mice, per condition. * P<0.05).

To further characterise the induced aggregates, we investigated the subcellular and regional location of tau lesions. We used recombinant tau assemblies to induce seeding owing to the high confidence that AT8-reactive aggregates result from seeded aggregation rather than the input tau. Within cell bodies, we observed large aggregates in peri-nuclear regions [Figure 4a]. Additionally, numerous smaller tau puncta were found along the length of neurites. Puncta were interrupted by regions apparently devoid of hyperphosphorylated tau. In contrast, Map2 staining revealed the presence of intact neurites, indicating that the punctate distribution of tau is not a consequence of neuronal fragmentation. They further demonstrate that neurons are able to tolerate tau aggregation to a certain degree without gross loss of morphology or overt toxicity. We compared levels of seeding between regions of the hippocampal slices. We observed the presence of AT8 positive structures in neurons within all subdivisions [Figure 4b]. However, AT8 reactivity was considerably greater within the CA1 region compared to CA2 and CA3. Approximately 80% of AT8-postitive structures were found in CA1, compared to ∼10% in each of CA2 and CA3 [Figure 4c]. We examined levels of tau as a potential underlying cause of CA1 susceptibility but observed comparable expression levels across different regions [Figure 4d]. In summary, these results demonstrate that challenge of OHSCs with assemblies of tau induces the accumulation of pathology in neurites and cell bodies, predominantly in CA1 neurons, resulting in widespread accumulation of intracellular hyperphosphorylated tau structures.

**Figure 4:**
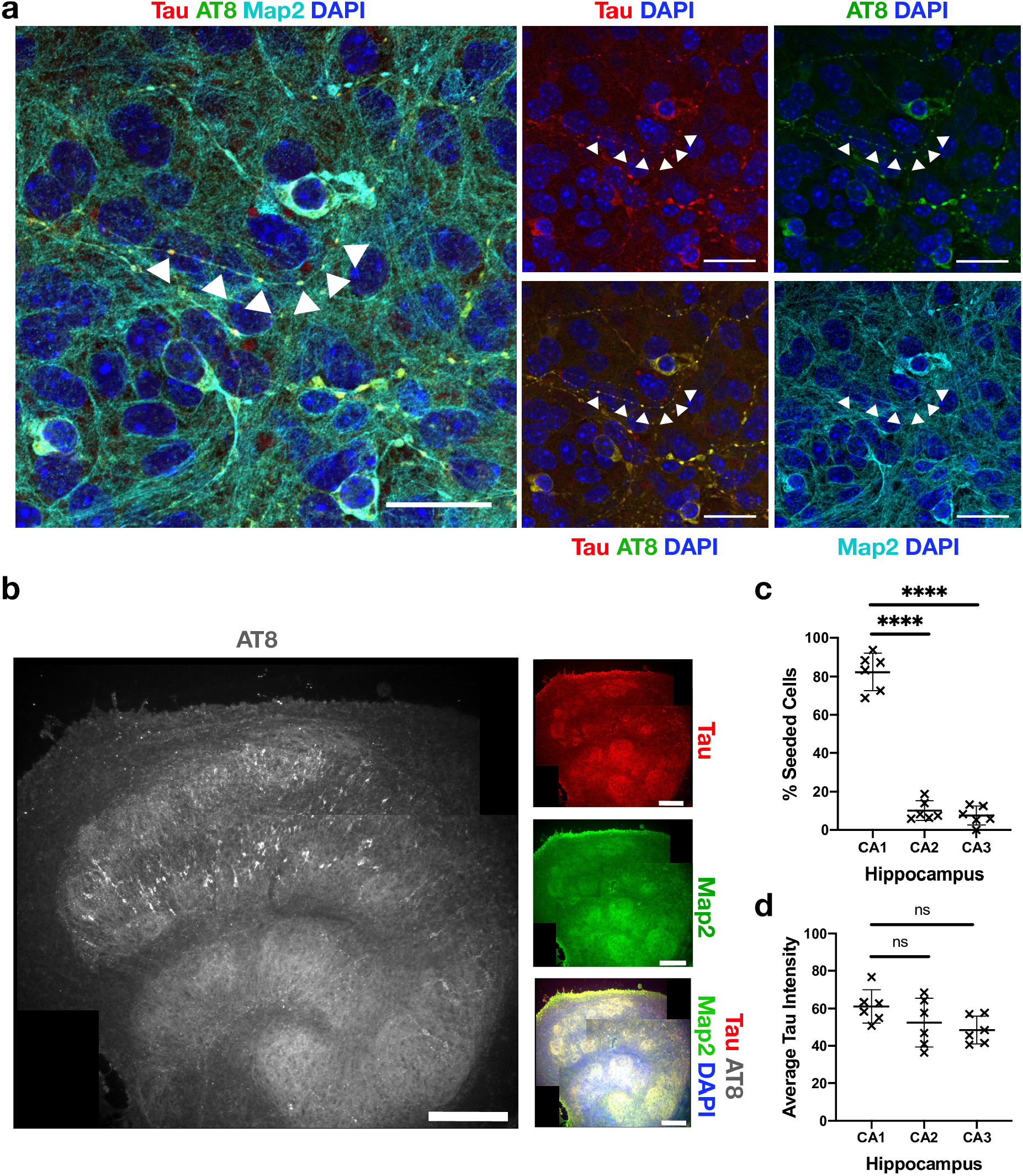
Neurons display phospho-tau aggregates within intact nerve processes and aggregates localise to CA1. **a**. OHSCs were challenged with 100 nM recombinant tau assemblies to induce seeded aggregation. Hyperphosphorylated tau puncta can be observed along intact nerve processes (arrows) and within cell bodies. Scale bars are 25 µm. **b**. Tiled image of representative OHSC challenged with 100 nM recombinant tau assemblies displays AT8 immunoreactivity predominantly in the CA1 subregion. Scale bars are 250 µm. **c**. The distribution of seeded cells in hippocampal subregions was quantified by counting cells positive for AT8 aggregates. **d**. Levels of tau, as quantified by pan-tau staining, show that CA1, CA2 and CA3 express similar levels of tau. Statistical significance determined by one-way ANOVA and Tukey’s post hoc multiple comparisons test (multiple fields imaged from slices from 6 different mice. **** P<0.0001).

To determine the time-dependence of this tau pathology, we next performed a time course following the addition of seed. Slices were fixed at 1, 2 or 3 weeks following challenge with 100 nM recombinant assemblies [Figure 5a]. Alternatively, slice cultures were fixed at 3 weeks following challenge with buffer only. As above, slices that were not exposed to tau assemblies developed no robust evidence of hyperphosphorylated tau puncta. However, challenge with tau assemblies resulted in increasing levels of bright AT8-positive structures over time, consistent with seeded aggregation of intracellular pools of tau [Figure 5b]. At 1 week after challenge, isolated AT8 positive puncta were observed as well as diffuse AT8 staining. A week later, puncta became more numerous and a few large aggregates were observed. However, 3 weeks after challenge with tau assemblies, AT8 staining was widespread with the presence of numerous aggregates that occupied entire cell bodies. The increase in AT8 staining followed an exponential curve with a doubling time of ∼7 days. The size of AT8-positive structures was similarly found to increase over time. Stained areas greater than 50 µm^2^, generally only present within cell bodies, were found to be largely absent at 1 week post-challenge but subsequently to increase in prevalence [Figure 5d]. This suggests that amplification of aggregates within individual neurons is driving the overall increase in AT8 signal. The results are therefore consistent with a model of growth of hyperphosphorylated tau structures via a process of templated aggregation following exposure to seed-competent tau assemblies.

**Figure 5:**
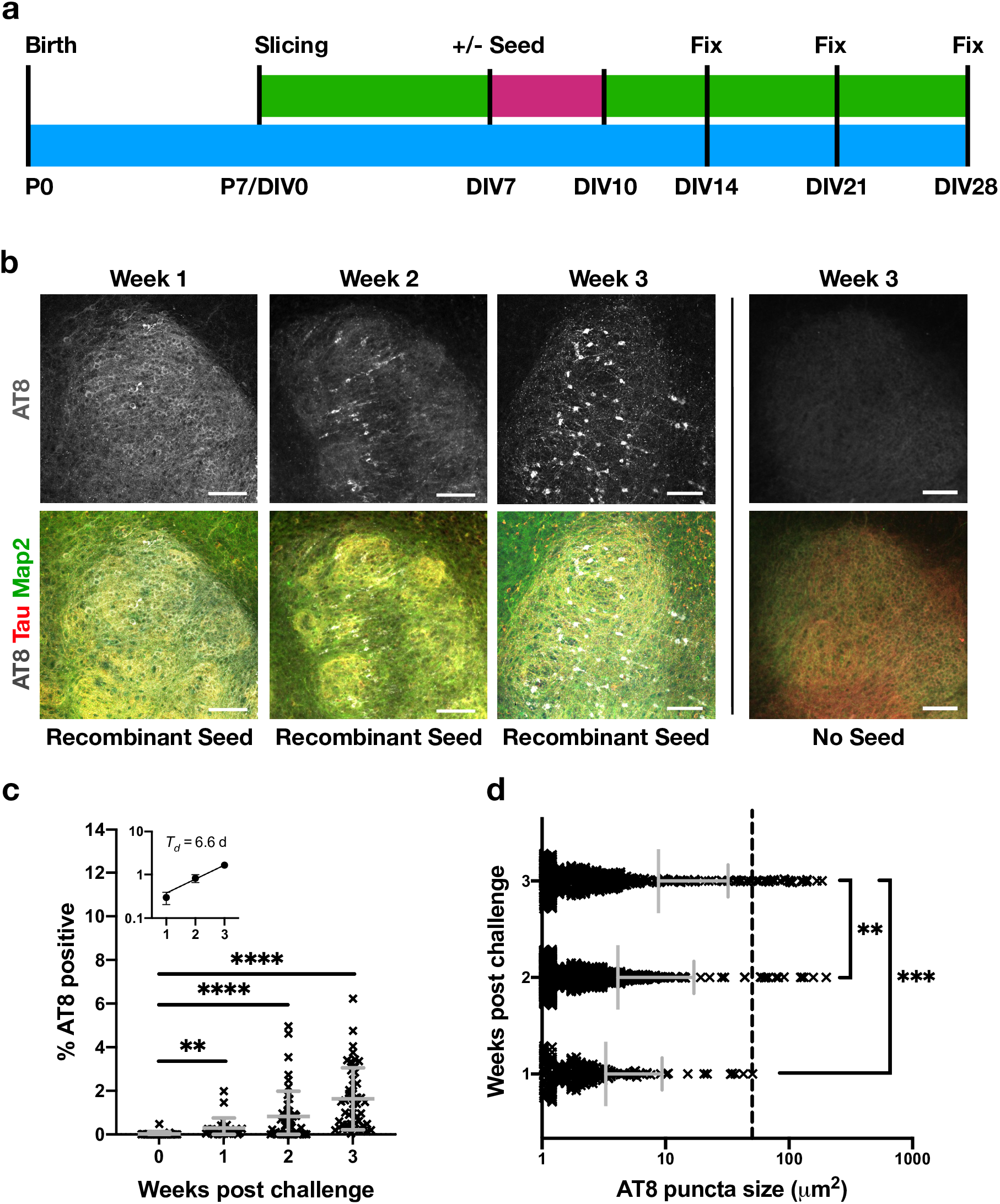
Phospho-tau reactivity and hyperphosphorylated tau aggregates increase over time. **a**. Schematic of OHSC preparation. 100 nM recombinant tau assemblies were added to the media as previous and left for 72 hours (pink) followed by a complete media change. Subsequently 50% media changes were performed twice weekly (green) until fixation at 1, 2 or 3 weeks post challenge. **b**. Slices fixed at 1 week post challenge display diffuse AT8 staining. Slices fixed at 2 or 3 weeks demonstrate increasing levels of puncta in cell bodies and neurites. OHSCs not challenged with exogenous tau fibrils exhibit only diffuse background levels of AT8 reactivity. Scale bars are 100 μm. **c**. Quantification of percent area that was AT8 reactive shows a significant increase in phospho-tau levels, with a doubling time of ∼7 days. Statistical significance determined by Kruskal-Wallis Test by ranks and Dunn’s multiple comparisons test (multiple fields imaged from slices from >2 different mice per time point. **P<0.01, **** P<0.0001). Inset represents the same data from weeks 1-3 plotted on a logarithmic scale. **d**. Quantification of AT8 positive puncta size shows an increase in the size of AT8 positive aggregates. Dotted line at 50 μm^2^ represents approximate lower size limit of cell body-occupying lesions. Statistical significance determined by Kruskal-Wallis Test by ranks and Dunn’s multiple comparisons test (multiple fields imaged from slices from >2 different mice per time point, **P<0.01, ***P<0.001).

The above results demonstrate that our OHSC model exhibits behaviour consistent with prion-like spread of tau. However, the dose we used (100 nM monomer equivalent) represents a concentration in excess of ISF and CSF tau concentrations, which occupy the low nanomolar to picomolar region. We therefore investigated the response of OHSCs to varying of the dose of exogenously-supplied tau assemblies. Remarkably, we found that a reduction of seed concentration from 100 nM to 30 nM resulted in virtually no seeded aggregation being detectable within the slice [Figure 6a]. Whereas cell bodies reactive for AT8 could be observed when challenged with 100 nM tau assemblies, only very rare and small AT8-positive assemblies in neurites were observed following challenge with 30 nM tau. Conversely, increasing exogenous tau concentration from 100 nM to 300 nM increased the AT8-immunorective area by almost 10-fold [Figure 6b]. To exclude any effect of the culture membrane on the efficiency of tau uptake, we applied tau at the same concentrations directly to the surface of the slices. We challenged OHSCs with 25 μl of recombinant tau aggregates applied to the apical surface of the slice. Alternatively, we supplied the same concentration of tau assemblies to the media as normal. Under both experimental set-ups we observed robust induction of seeding at 100 nM, but not at 30 nM [Figure 6c]. Thus, the local concentration of tau governs seeded aggregation and is independent of application route. These results demonstrate that tau seeding in OHSCs only occurs efficiently at concentrations above 100 nM of supplied assemblies.

**Figure 6:**
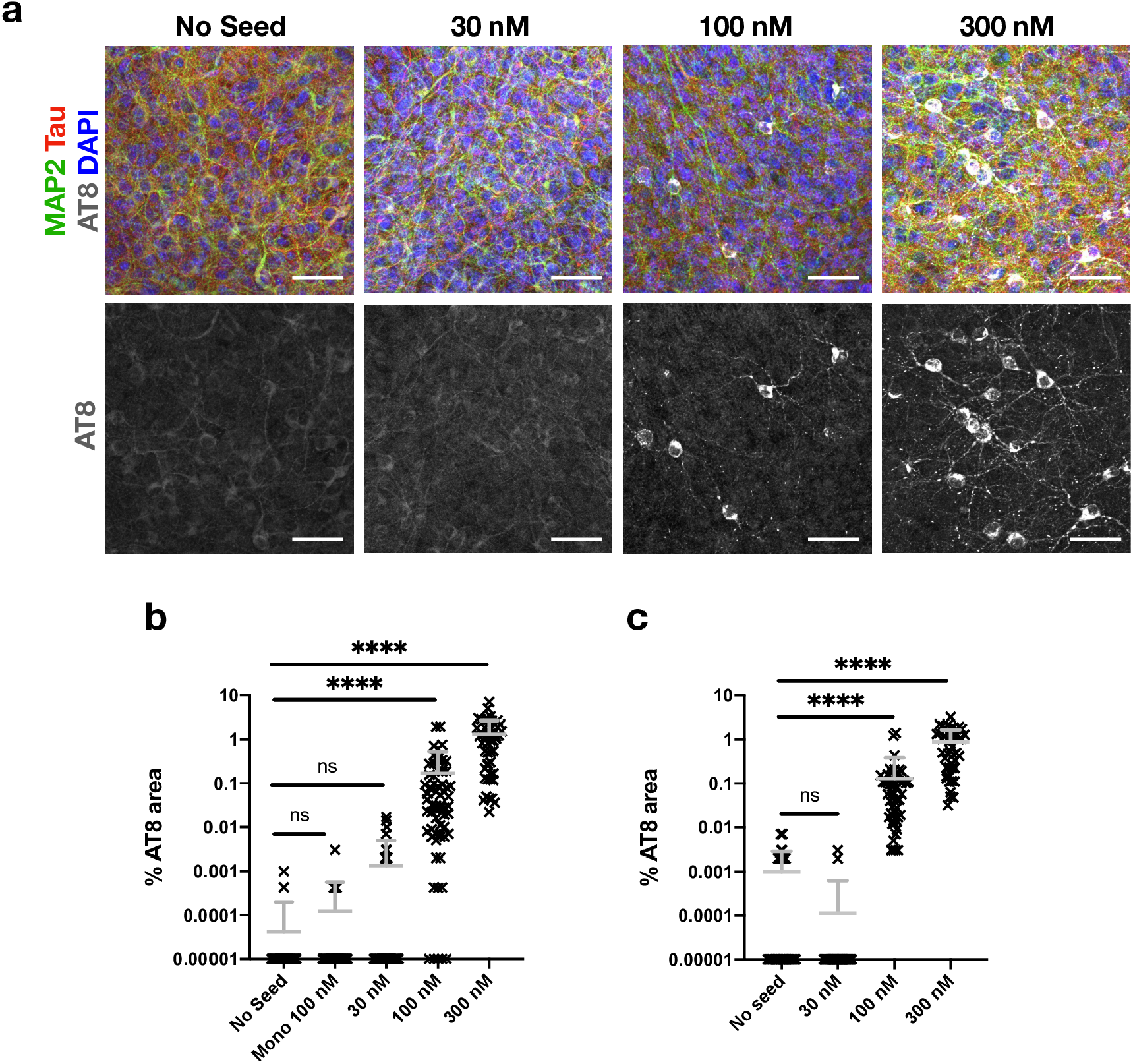
Seeding of tau occurs with an apparent threshold. **a**. OHSCs challenged with 30 nM, 100 nM and 300 nM of recombinant tau assemblies, or with buffer only. Scale bars are 50 μm. **b**. Quantification of seeding levels in P301S OHSCs, upon the addition of 30 nM, 100 nM or 300 nM recombinant tau assemblies, 100 nM tau monomer or buffer only underneath the culture insert. Statistical significance determined by Kruskal-Wallis Test by ranks and Dunn’s multiple comparisons test (Slices from 3 different mice, per condition. **** P<0.0001). **c**. Quantification of seeding levels in P301S OHSCs, upon the addition of recombinant tau assemblies or buffer only to the apical surface of individual slices. Statistical significance determined by Kruskal-Wallis Test by ranks and Dunn’s multiple comparisons test (Slices from 3 different mice, per condition. **** P<0.0001).

Independently acting infectious particles such as viruses retain infectivity upon dilution until they are diluted out at endpoint. They display one-hit dynamics where proportion of infected cells, P(I), can be described by the equation P(I) = 1 *– e*^*-m*^ where m is the average number of infectious agents added per cell. To determine whether tau assemblies display these properties, we titrated tau assemblies on HEK293’s expressing tau-venus. Here, where conditions have been optimised for sensitive detection of seeding, and tau assemblies are delivered directly to the cytoplasm with transfection reagents, we observe that seeding activity is proportional to dose and can be titrated down. The observed level of seeding approximates a one-hit titration curve [Figure 7a]. Thus, tau assemblies have the intrinsic ability to act as independent particles when tested in reporter cell lines. This is in direct contrast to the results observed in OHSCs where seeding reduces much more rapidly as tau assemblies are diluted than would be expected under a single-hit model [Figure 7a]. One potential explanation for these differences is that clearance mechanisms in intact tissue inherently prevent single-particle activity. To test this, we titrated AAV1/2.hSyn-GFP particles expressing GFP and measured the percent of Map2-positive neurons that were transduced. We observed that AAV1/2 behaved in a manner consistent with one-hit dynamics [Figure 7b]. Thus, tau assemblies differ from classical infectious agents and do not titrate in a manner expected of independently acting particles in mouse neural tissue. Rather, seeding is a behaviour that only emerges at high concentrations of extracellular tau assemblies.

**Figure 7:**
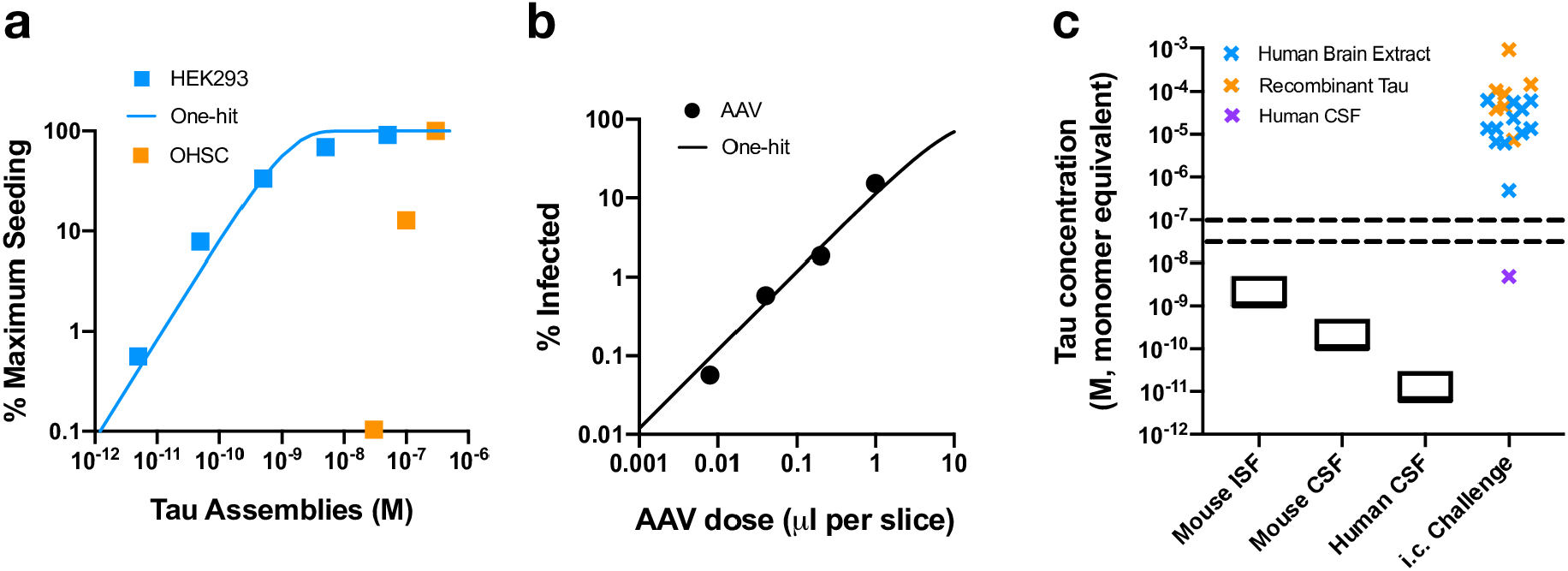
Tau seeding does not conform to one-hit dynamics. **a**. Tau assemblies were titrated on HEK293 tau-venus cells using LF2000 transfection reagent, and the amount of seeding was quantified and expressed as percent of maximum. A one-hit curve was fitted using values outside the plateau. Tau seeding in OHSCs, means derived from Fig 6b, cannot be fitted to a one-hit model. **b**. Infection of P301S OHSCs with AAV1/2.hSyn-GFP with one-hit curve fitted to all data points (Slices from 3 different mice, per condition). **c**. Comparison of the concentration of tau used in stereotaxic injection experiments, coloured by origin, with ranges of ISF and CSF concentrations of tau measured in mice and humans, sourced from the literature (see Supp Table 1). The dotted lines represent the apparent threshold for the seeded aggregation of tau in neural tissue observed here at between 30-100 nM tau.

## Discussion

Elucidating the mechanism of tau aggregation and its apparent spread through the brain is critical to the development of mechanism-based therapeutics. The ‘prion-like’ model of tau spread posits that the transit of assembled tau species from affected to naïve cells promotes the exponential spread of pathological tau over time and space within a diseased brain. In support of this model, extracellular fluids of tauopathy patients’ brains contain seed-competent tau species: CSF samples from both AD and Pick’s disease patients give rise to seeded aggregation in biosensor cell lines and biochemical detection assays [26–28]. Further evidence in support of the prion-like model comes from *in vivo* challenge experiments: intracranial injection of assembled tau can result in induced tau pathology in wildtype or tau-transgenic rodent brains. Understanding how these *in vivo* challenge experiments, which are typically performed at high concentration [Figure 7c, Supplementary Table 1], translate to physiological concentrations is important in order to assess the applicability of these results to disease mechanisms. Contrary to our expectations, we found that tau seeding activity rapidly dropped away upon dilution in OHSCs. Observations of seeding behaviour at high concentration therefore cannot necessarily be extrapolated down to inform on the behaviour of tau at lower concentrations.

In reporter cells, we observed that tau seeds titrated in a one-hit manner, as expected of independently acting particles. This suggests that reporter cell lines which have been validated in this way can be used to ascertain the intrinsic seeding activity of tau preparations, which can then be expressed as seeding units per quantity tau, analogous to other infectious agents. In OHSCs, tau seeding was observed at concentrations in excess of 100 nM. Further dilution of tau assemblies prevented seeded aggregation long before end-point dilution of seeds therefore displaying a marked deviation from one-hit dynamics. Such deviations in virus infectivity can be caused by host cell factors that prevent infection becoming saturated by high viral dose [29,30]. By analogy, we consider it likely that homeostatic mechanisms act to prevent seeded aggregation of tau but become saturated by high tau concentrations. The nature of any such saturable barrier to seeding is not clear. One possibility is that phagocytic cells present in slices preclude observations of one-hit dynamics. This was not the case, however, since AAV particles were found to titrate with one-hit dynamics in OHSCs. A trivial explanation of tau at low concentration being unable to cross the membrane was also ruled out. Other mechanisms are therefore implicated such as saturation of proteostatic mechanisms or uptake to the cell. Identification of these defences may provide a valuable route to understanding the mechanisms which prevent prion-like propagation, and their potential deterioration in disease.

Our results suggest that healthy neural tissue is able to withstand the concentration of tau present in extracellular fluids without observable seeded aggregation. The effective threshold for seeding, measured here at around 100 nM, exceeds physiological ISF/CSF concentrations by several orders of magnitude. Thus, in order for spreading via seeded aggregation to occur, our results suggest that other mechanisms are required. For instance, uncontrolled neuronal cell death or release of tau into synaptic clefts may transiently raise the local concentration of tau to high levels. Alternatively, the threshold for seeded tau aggregation may be altered in the degenerating brain, for instance through inflammation or other mechanisms. Finally, other modes of transmission within the brain that do not rely on naked pools of extracellular tau may circumvent the non-linear dose response observed here. Such mechanisms include tau spreading via tunnelling nanotubules and in extracellular vesicles [31,32].

The tau species present in the extracellular spaces of the brain are likely to differ from those used here in terms of pathological fold, post-translational modification and proteolytic truncation. Potentially of interest in this regard is the study by Skachokova and colleagues who successfully induced seeding following injection of P301S tau transgenic mice with a 1,000-fold concentrate of CSF from AD patients [33]. At 5-17 nM, these samples are still far in excess of human CSF tau concentrations. But, notably, these concentrations are below the threshold defined in our OHSCs model [Figure 7c]. Future work is therefore required to determine whether mechanisms not captured here but present within the degenerating brain enable seeded aggregation to occur. Future studies should therefore seek to develop seeding in wildtype, preferably human, settings in order to assess the nature of seeding of human brain-origin tau assemblies.

Our experiments demonstrated that neurons in CA1 were particularly sensitive to seeded aggregation compared to those in CA2 and CA3. Whilst injection of tau assemblies to the *in vivo* brain also demonstrates prominent CA1 seeding, proximity to the injection site is the major determinant of seeding in animal studies, thereby confounding conclusions of regional susceptibility [3,5,34]. We found that tau substrate levels were not implicated in the phenotype, suggesting that other factors are responsible for the increased susceptibility. These results are potentially of interest in the study of selective vulnerability since it is well established that CA1 displays more pronounced AT8 reactivity in post-mortem human brains [8,35,36]. In humans, the advanced pathology in CA1 versus other HC regions could potentially be explained either by selective vulnerability of its neurons to aggregation, or by its upstream position in the circuitry of the HC and therefore prone to earlier and more pronounced pathology under a spreading model. Our findings lend support to an underlying increased susceptibility of CA1 neurons to pathology. OHSCs therefore provide a suitable platform for future studies to determine the biological basis of this susceptibility.

Our findings help define the prion-like characteristics of tau assemblies. Whilst intrinsic seeding activity that titrates according to one-hit dose-response can be detected in biosensor assays, this behaviour is lost in neural tissue. Our findings suggest that neural tissue possesses homeostatic mechanisms that are capable of successfully preventing seeded aggregation. Saturating levels of tau assemblies are required to overcome these barriers to initiate seeded aggregation.

## Materials and Methods

### Mouse lines

All animal work was licensed under the UK Animals (Scientific Procedures) Act 1986 and approved by the Medical Research Council Animal Welfare and Ethical Review Body. P301S tau transgenic mice [21] that had been extensively backcrossed to C57BL/6 background were obtained from Dr Michel Goedert, MRC Laboratory of Molecular Biology, UK. Male and female were used in the study and humanely sacrificed by cervical dislocation.

### Recombinant tau production

The expression and purification of recombinant human 0N4R tau bearing the P301S mutation from *E. coli* BL-21 (DE3, Agilent Technologies) was performed as described previously (Goedert and Jakes, 1990) with small modifications. Bacterial pellets were collected through centrifugation (3300 g, 4 °C, 10 min) and then resuspended in 10 ml/l of culture with buffer A (50 mM MES pH 6.5, 10 mM EDTA, 14 mM β-mercaptoethanol, 0.1 mM PMSF, 1 mM benzamidine, 1x complete EDTA-free protease inhibitors). The resuspended bacteria were lysed on ice using a probe sonicator (approximately 60% amplitude) and then boiled for 10 min at 95 °C to pellet the majority of proteins, while tau will remain in solution as a natively unfolded protein. Denatured proteins were pelleted through ultracentrifugation (100,000 g, 4 °C, 50 min). The clarified supernatant containing monomeric tau P301S was then passed through a HiTrap CaptoS (Cytiva) cation exchange column and the bound proteins were eluted through a 0-50 % gradient elution with Buffer A containing 1 M NaCl. Eluted fractions were assessed through SDS-PAGE and total protein staining with Coomassie InstantBlue. Fractions of interest were concentrated using 10 kDa cut-off Amicon Ultra-4 concentrators (Merck Millipore) before loading on a Superdex 200 10/300 GL (Cytiva) size exclusion chromatography column. The final tau P301S protein was stored in PBS containing 1 mM DTT. All the affinity purification and size exclusion chromatography steps were performed using the ÄKTA Pure system (Cytiva).

### Recombinant Tau Aggregation

Tau monomer was added to aggregation buffer (20 μM Heparin, 60 μM P301S tau monomer, 1x complete EDTA-free protease inhibitors, 2 µM DTT in PBS) and incubated at 37 °C for 3 days. The resulting P301S tau filaments were sonicated for 15 seconds before long-term storage at −80 °C.

### ThioflavinT Assay

Tau monomer was added to aggregation buffer, with 10 µM sterile filtered ThioflavinT (ThT). Samples were loaded in triplicate into black 96-well plates. Plates were loaded into a CLARIOstar (BMG Labtech), and measurements were taken every 5 minutes after shaking, for 72 hours at 37 °C min (excitation and emission wavelength 440 nm and 510 nm respectively).

### TEM

Recombinant tau fibrils were mounted on carbon-coated copper grids (EM Resolutions) via suspension of the grid on a single droplet. The grid was then stained with 1% uranyl acetate and imaged with a FEI Tecnai G20 electron microscope operating at 200kV and an AMT camera.

### Preparation of tau assemblies from brains and OHSCs

Tau was extracted from aged brains (26 weeks) from mice transgenic for human P301S tau using sarkosyl extraction. Tissues were homogenised for 30 s in 4 volumes of ice-cold H-Buffer (10mM Tris pH 7.4, 1mM EGTA, 0.8M NaCl, 10% sucrose, protease and phosphatase inhibitors (Halt™ Protease and Phosphatase Inhibitor Cocktail)) using the VelociRuptor V2 Microtube Homogeniser (Scientific Laboratory Supplies). The homogenates were spun for 20 minutes at 20,000 × g and supernatant was collected. The resulting pellet was re-homogenised as above in 2 volumes of ice-cold H-Buffer and processed as above. Supernatants from both spins were combined and sarkosyl was added to a final concentration of 1% and incubated for 1 h at 37 °C. Supernatants were then spun at 100,000 × g at 4 °C for 1 h. The resulting pellet was resuspended in 0.2 volumes of PBS and sonicated for 15 s in a water-bath sonicator before storage at −80 °C. For OHSCs, the same procedure was followed, except slices were freeze thawed 5 times in 20 μl per slice ice-cold H-Buffer and the final pellet was resuspended in 5 μl per slice PBS.

### Western blotting

Samples were transferred to fresh microcentrifuge tubes, to which appropriate volumes of 4× NuPAGE LDS sample buffer (Thermo Fisher) containing 50 mM DTT was added and heated to 95 °C for 5 min. Samples were resolved using NuPAGE Bis–Tris Novex 4–12% gels (Life Technologies) and electroblotted to a 0.2-μm PVDF membrane using the Transblot Turbo Transfer System (Bio-Rad). Membranes were blocked with 5% milk TBS–Tween 20 before incubation with primary antibodies overnight at 4 °C. Membranes were then probed with appropriate secondary antibodies conjugated with HRP for 1 h. Membranes were washed repeatedly in TBS–0.1% Tween-20 after both primary and secondary antibody incubation. Blots were incubated with Pierce Super Signal or Millipore Immobilon enhanced chemiluminescence reagents for 5 min and visualised using a ChemiDoc system (Bio-Rad).

### Dot Blot

Recombinant or mouse-extracted tau fibrils were diluted in PBS as indicated in Supplementary Figure 1 and applied to 0.2 µm nitrocellulose membrane using the Bio-Dot microfiltration apparatus (Bio-Rad). The membranes were then blocked in 5% milk TBS-Tween 20 and subsequently incubated with primary antibody overnight. The next day, the membranes were probed with appropriate secondary antibodies conjugated with Alexa488 fluorophore and imaged using the ChemiDoc system (Bio-Rad). The dot intensities were quantified with the Image Studio Lite software (LI-COR Biosciences) and the values for the recombinant fibrils were fitted to a simple linear regression curve.

### Seeding assay in HEK293

The seeding assay was carried out as described previously [25]. Briefly, HEK293 P301S tau-venus cells were plated at 15,000 cells per well in black 96-well plates pre-coated with poly D-lysine in 50 µL OptiMEM (Thermo Fisher). Tau assemblies were diluted in 50 µL OptiMEM (Thermo Fisher) and added to cells with 0.5 µl per well Lipofectamine 2000. After 1.5 h, 100 µL complete DMEM was added to each well to stop the transfection process. Cells were incubated at 37 °C in an IncuCyte® S3 Live-Cell Analysis System for 48 − 72 h after addition of fibrils.

### Preparation and culturing of organotypic slices

Organotypic hippocampal slice cultures were prepared and cultured according to the protocols described previously [19,23]. Brains from P6-P9 pups were rapidly removed and kept in ice-cold Slicing medium (EBSS + 25 mM HEPES+ 1x Penicillin/Streptomycin) on ice. All equipment was kept ice-cold. Brains were bisected along the midline and the cerebellum was removed using a sterile scalpel. The medial, cut surface of the brain was adhered to the stage of a Leica VT1200S Vibratome using cyanoacrylate (Loctite Super Glue) and the vibratome stage was flooded with ice-cold Slicing medium. Hemispheres were arranged such that the vibratome blade sliced in a rostral to caudal direction. Sagittal slices of 300 µm thickness were prepared and the hippocampus was sub-dissected using sterile needles. Hippocampal slices were transferred to 15 mL tubes filled with ice-cold Slicing medium using sterile plastic pipettes with the ends cut off. Slices were then transferred onto sterile 0.4 μm pore membranes (Millipore PICM0RG50) in 6-well plates pre-filled with 1 mL pre-warmed Culture medium (50% MEM with GlutaMAX, 18% EBSS, 6% EBSS+D-Glucose, 1% Penicillin-Streptomycin, 0.06% nystatin and 25% Horse Serum) and incubated at 37 °C in a humid atmosphere with 5% CO2. Three slices were typically maintained per well. 24 h after plating 100% media was exchanged and thereafter a 50% media exchange was carried out twice per week. For seeding experiments, tau assemblies were diluted in Culture medium and added to the underside of the membrane with 100% media change. After three days, assemblies were removed by 100% media change.

### Adeno-associated virus

AAV1/2.hSyn-GFP particles were generated by co-transfection of HEK293T cells with AAV2/1 (Addgene 112862), AAV2/2 (Addgene 104963), adenovirus helper plasmid pAdDeltaF6 (Addgene 112867) and pAAV-hSyn-EGFP (Addgene 50465). Virus particles were purified by iodixanol gradient in at T70i ultracentrifuge rotor as previous [37]. Viral purity was confirmed by the presence of three bands following SDS-PAGE and staining with Coomassie InstantBlue.

### Immunofluorescence microscopy

Slices on membranes were washed with PBS and then fixed in 4% (w/v) paraformaldehyde for 20 min at 37 °C. Subsequently, membranes were rinsed 2–3 times with PBS and left shaking gently for 15 min to remove traces of paraformaldehyde before subsequent processing. Slices were permeabilised with 0.5% (v/v) Triton X-100 in immunofluorescence blocking buffer (IF block) (3% goat serum in 1× PBS) for 1 h at room temperature, and rinsed with 3x with TBS. Slices were then incubated with primary antibodies diluted in IF block overnight at 4 °C, rinsed with 3 times with TBS, and incubated for 2 h in the dark with secondary antibodies, also diluted in IF block. Secondary antibodies conjugated to Alexa Fluor 488, 568 or 647 were obtained from Thermo Fisher. Following rinsing 3x with TBS, the slices were incubated with Hoechst stain for 10 min and rinsed 3x with TBS. Membranes were placed on slides (slice side up), mounting medium (ProLong Diamond, Life Technologies) was added and a cover slip was placed on top of the slice. Images were captured using a Zeiss LSM780 Confocal Microscope with either a 20x or a 63x objective lens. Images were collected and stitched, where appropriate, using ZEISS Zen software package.

### Image Analysis and statistics

For tau seeding assays in HEK293 cells, aggregates were detected and quantified using the ComDet plugin in Fiji [38]. Threshold levels for detection of aggregates were adjusted using mock-seeded images for each experiment. Levels of seeding were calculated as (number of aggregates)/(total cells) × 100 for individual fields. For slice cultures, maximum intensity Z-projections were interrogated for AT8 immunoreactivity by the application of a binary threshold-based mask in ImageJ. Percent area of AT8 reactivity was determined in regions of 100 x 100 μm. GFP positive neurons upon AAV infection were analysed in the same way. For measures of number of neurons affected in hippocampal subregions, a manual count of cell bodies positive for AT8 immunoreactivity was performed. Zero values were given an arbitrary value of 10^−5^ for representation on log-scale axes. The data in all graphs are represented as the mean +/- SD. Data was analysed via the Kruskal-Wallis test by ranks, unless it was determined to be normally distributed, in which case a one-way ANOVA was employed. All statistics were carried out in GraphPad Prism Version 8.

## Acknowledgments

We thank Dr Michel Goedert for provision of P301S tau transgenic mice and Cambridge Advanced Imaging Centre for access to electron microscope facilities. We thank Prof Sir David Klenerman for valuable discussions on the early versions of this manuscript.

## Author contributions

Conceived research: WAM, MPC, ASM, LVCM; Designed experiments: ASM, LVCM, WAM; Developed slice culture assay: ASM, LVCM, CD, OS, CK, MJV, LCJ; Performed experiments and analysed data: ASM, LVCM, TK, BJT, WAM, SS, SC. All authors contributed to writing and editing of the manuscript.

## Ethics Approval

All animal work was licensed under the UK Animals (Scientific Procedures) Act 1986 and approved by the Medical Research Council Animal Welfare and Ethical Review Body.

