## Supplementary data for "Tau assemblies do not behave like independently acting prion-like particles in mouse neural tissue"

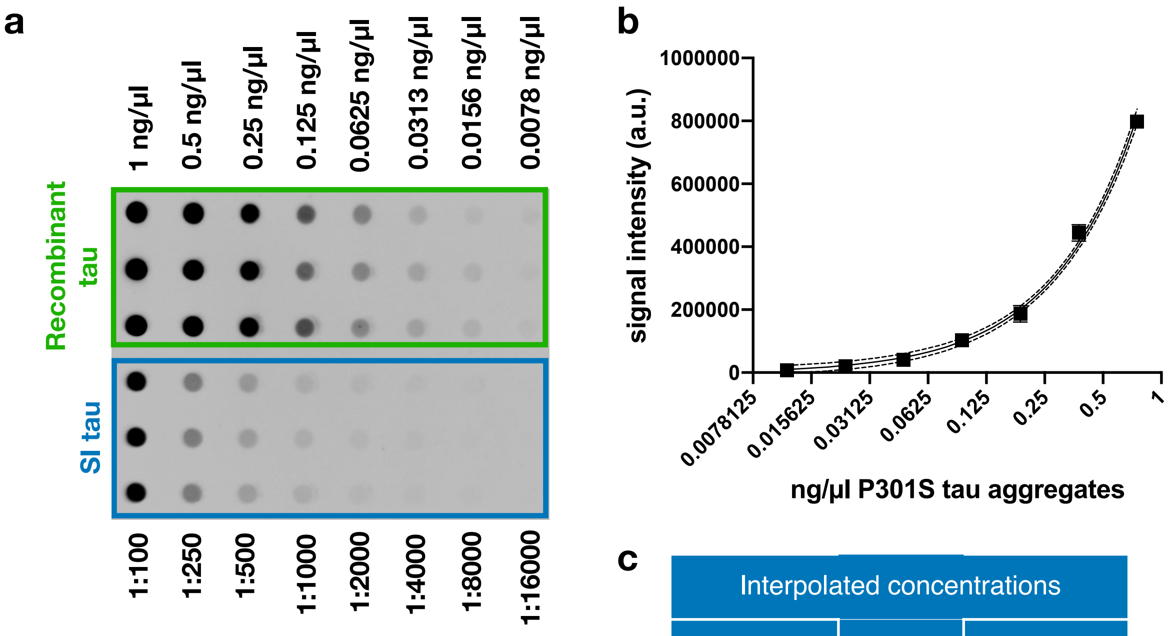


c

| Interpolated concentrations | | |
|---|---|---|
| | (mg/ml) | µM |
| SI tau | 2.85 | 71 |

**Supplementary Figure 1: Quantification of SI mouse brain derived tau**

**a.** A dot blot titration of SI mouse brain derived tau, with recombinant tau assemblies at the indicated concentrations used as a standard. **b**. Standard curve generated from dot blot signal intensities. **c.** Interpolated concentration of SI mouse brain derived tau.

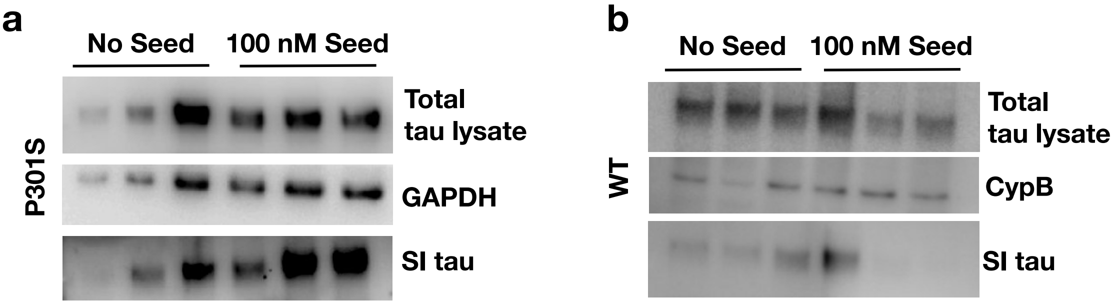


**Supplementary Figure 2: SI extraction of P301S and WT OHSCs**

**a.** Western blot of the total tau in P301S OHSC lysate and the SI fraction of P301S OHSCs with and without the addition of 100 nM recombinant tau assemblies. **b.** Western blot of the total tau in WT OHSC lysate and the SI fraction of WT OHSCs with and without the addition of 100 nM recombinant tau assemblies.

| Mouse Model | Tau Source | Concentration | Molar Concentration | Reference |
|---|---|---|---|---|
| PS19 | K18 Recombinant Tau | 2 mg/ml | 145 µM | [1] |
| PS19 | P301S Recombinant Tau | 2 mg/ml | 44 µM | [1] |
| P301L | K18 Recombinant Tau | 12.5 mg/ml | 912 µM | [2] |
| PS19 | AD Brain Extract | 2.5 mg/ml | 50 µM | [3] |
| PS19 | CBD Brain Extract | 20 µg/ml | 400 nM | [3] |
| WT | AD Brain Extract | 1.6 mg/ml | 38 µM | [4] |
| WT | WT Recombinant Tau | 1.8 mg/ml | 39 µM | [4] |
| PS19 | P301S Recombinant Tau | 4 mg/ml | 87 µM | [5] |
| WT | AD Brain Extract | 2.56 mg/ml | 51 µM | [6] |
| WT | CBD Brain Extract | 0.256 mg/ml | 5.1 µM | [6] |
| WT | PSP Brain Extract | 0.44 mg/ml | 8 µM | [6] |
| WT and FAD | AD Brain Extract | 1 mg/ml | 20 µM | [7] |
| P301S | P301S Recombinant Tau | 0.31 mg/ml | 7.3 µM | [8] |
| hTau | AD Brain Extract | 0.275 mg/ml | 6.6 µM | [9] |
| P301S | AD CSF | 204.6 ng/ml | 4.08 nM | [10] |
| 6hTau | AD Brain Extract | 2.3 mg/ml | 56 µM | [11] |
| 6hTau | PSP Brain Extract | 0.56 mg/ml | 14 µM | [11] |
| 6hTau | CBD Brain Extract | 0.56 mg/ml | 14 µM | [11] |
| 6hTau | PiD Brain Extract | 0.56 mg/ml | 14 µM | [11] |
| WT | WT Recombinant Tau | 4.5 mg/ml | 90 µM | [12] |

**Supplementary Table 1: Summary of reported concentrations of tau used in stereotaxic injection experiments showing seeded aggregation of tau.**

Stereotaxic injection experiments separated by the source of tau used. Molar concentrations calculated from the types of tau used. The average molecular weight of the six tau isoforms was used in the case of human brain extract.
